# Telescoping bimodal latent Dirichlet allocation to identify expression QTLs across tissues

**DOI:** 10.1101/2021.10.27.466156

**Authors:** Ariel DH Gewirtz, F William Townes, Barbara E Engelhardt

**Affiliations:** Lewis-Sigler Institute of Integrative Genomics, Princeton University, Washington Road, Princeton, USA; Department of Computer Science, Princeton University, 35 Olden Street, 08540 Princeton, USA; Gladstone Institutes, 1700 Owens Street, 94158 San Francisco, USA

**Keywords:** eQTLs, multi-omic data, latent variable models

## Abstract

Expression quantitative trait loci (eQTLs), or single nucleotide polymorphisms (SNPs) that affect average gene expression levels, provide important insights into context-specific gene regulation. Classic eQTL analyses use one-to-one association tests, which test gene-variant pairs individually and ignore correlations induced by gene regulatory networks and linkage disequilibrium. Probabilistic topic models, such as latent Dirichlet allocation, estimate latent topics for a collection of count observations. Prior multi-modal frameworks that bridge genotype and expression data assume matched sample numbers between modalities. However, many data sets have a nested structure where one individual has several associated gene expression samples and a single germline genotype vector. Here, we build a telescoping bimodal latent Dirichlet allocation (TBLDA) framework to learn shared topics across gene expression and genotype data that allows multiple RNA-sequencing samples to correspond to a single individual’s genotype. By using raw count data, our model avoids possible adulteration via normalization procedures. Ancestral structure is captured in a genotype-specific latent space, effectively removing it from shared components. Using GTEx v8 expression data across ten tissues and genotype data, we show that the estimated topics capture meaningful and robust biological signal in both modalities, and identify associations within and across tissue types. We identify 53,358 cis-eQTLs and 1,173 trans-eQTLs by conducting eQTL mapping between the most informative features in each topic. Our TBLDA model is able to identify associations using raw sequencing count data when the samples in two separate data modalities are matched one-to-many, as is often the case in biological data.

## Background

Genomic differences, such as single nucleotide polymorphisms (SNPs), among individuals are important drivers of gene expression variability. Much previous work has focused on discovering expression quantitative trait loci (eQTLs), which capture associations between the number of copies of a minor allele present at a given genomic locus and the expression level of a single gene (GTEx Consortium 2017; GTEx Consortium et al. 2020). However, a one-to-one mapping of genes to SNPs is too simplistic given the reality of biological interactions and the availability of many observations per individual. Pleiotropy, gene regulatory networks with biological redundancy and feedback loops, and linkage disequilibrium (LD) blocks of highly-correlated SNPs all contribute to a complex and dynamic biological regulatory system.

From a statistical perspective, performing genome-wide one-to-one association tests yields an astronomical multiple testing burden for trans-eQTLs, where the agnostic approach examines every interchromosomal gene and SNP combination. Statistical power is further reduced because trans-eQTLs, or eQTLs where the regulatory SNP is on a different chromosome than the gene that it regulates, often have smaller effect sizes than cis-eQTLs, or eQTLs where the regulatory SNP is local to the gene that it regulates (Petretto et al. 2006). One method to reduce the effective number of tests is to cluster correlated SNPs and genes, and compare the averaged cluster signals, versus testing for every possible marginal association.

Probabilistic topic models, such as latent Dirichlet allocation (LDA), are unsupervised machine learning methods that were initially introduced in natural language processing (Blei et al. 2003) and in statistical genetics as models of ancestry (Pritchard et al. 2000). LDA finds latent topics via soft clustering of feature counts over many samples while simultaneously estimating each sample’s topic membership proportions. More recently, these types of models have been applied to gene expression data with gene counts as features. The topics estimated by these models represent interpretable underlying biology such as cell type or developmental stage, and have been used in QTL mapping as the quantitative traits themselves (Dey et al. 2017; Hore et al. 2016).

We hypothesized that multi-modal topic modeling could identify clusters of co-regulated genes and SNPs. Existing methods have used Dirichlet process mixture models to integrate two data modalities (Savage et al. 2010), but nonparametric Bayesian models tend to be too computationally intense for larger data sets such as modern genotype arrays, which capture millions of SNPs. Argelaguet et al. (2018) designed a factor model framework (MOFA) to jointly model multiple data modalities, allowing various data likelihoods via link functions. However, relevant methods assume that the modalities are measured on the same samples such that there is a single observation from each individual in each modality (Argelaguet et al. 2018; Ash et al. 2021; Li and Gaynanova 2018; Savage et al. 2010; Virtanen et al. 2012; Zhao et al. 2016). Many earlier methods also require gene expression data to be normalized, potentially adulterating true signals or spuriously adding false ones (Hicks et al. 2018; Hore et al. 2016; Love et al. 2014; Robinson et al. 2010).

In this work, we create a probabilistic model to find shared structure between gene expression and genotype data. Our model uses raw sequencing read counts and works with a nested data structure where, although samples are paired, modalities may have different numbers of samples from each subject. This is often the case when we have many samples of gene expression from a particular donor—as in the GTEx data with multiple tissue samples per donor, and also for single-cell RNA-sequencing samples with multiple cells per subject—but a single germline genotype vector. We apply our model to GTEx v8 data, and use known sample tissue labels and cell type enrichment scores to interpret the biological context of the estimated components (GTEx Consortium et al. 2020). To demonstrate the model’s ability to find shared variation between data modalities, we conduct eQTL mapping using the most informative features in each topic to find both known and novel—and tissue-specific and general— cis- and trans-eQTLs.

## Methods

Given a genotype matrix and an RNA-sequencing expression matrix, our goal is to find latent factors that capture groups of SNPs and genes that covary across samples. Assume that we have two matrices as inputs: a RNA-seq count matrix 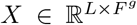 for *F^g^* genes across *L* samples and a genotype matrix 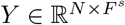 in minor allele dosage format (0,1, 2) for *F^s^* SNPs across *N* individuals. Our model assumes that there are *K* latent topics where (i) each sample ℓ ∈ {1, …, *L*} has membership proportion *ϕ*_*i*ℓ*k*_, (ii) each individual *i* ∈ {1,…, *N*} has membership proportion *θ_ik_* such that 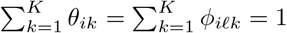, and (iii) each sample has a known mapping to exactly one of the individuals (Fig. 1, S1). Topics are modeled as distributions over features, where, similar to LDA, gene expression topics λ*^g^* are located on the simplex (Blei et al. 2003), while genotype topics λ*^s^* are modeled independently over SNPs *j* ∈ {1,…, *F^s^*} as in the Structure model (Pritchard et al. 2000). The formal model is as follows:

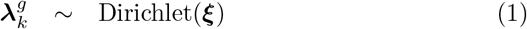

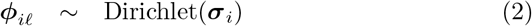

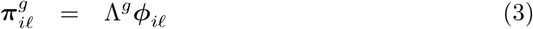

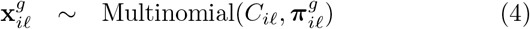

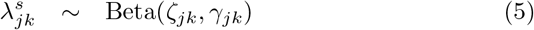

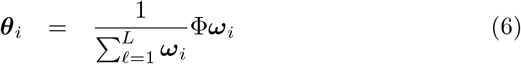

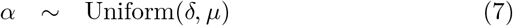

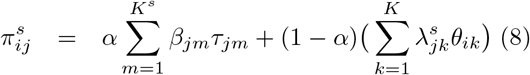

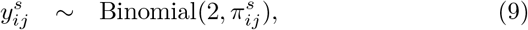

where *C_l_* is the total gene count in sample ℓ, and 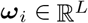 is an indicator vector for individual *i*, where *ω*_*i*ℓ_ = 1 if sample *l* originates from individual *i*. Here, we denote the matrices Φ, Λ^*s*^, and Λ^*g*^ as concatenated column vectors ***ϕ*_*i*ℓ_**, 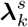, and 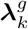. We use stochastic variational inference to compute posterior estimates for Φ, Λ^*s*^, and Λ*^*g*^* (see Methods for details).

**Figure 1.**
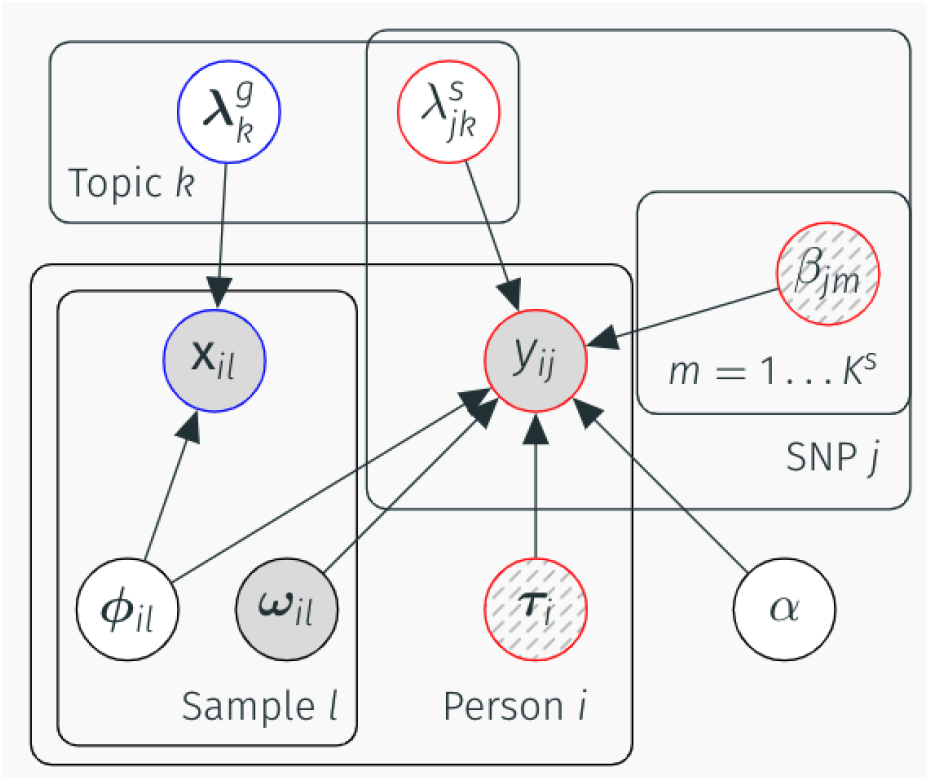
TBLDA model plate diagram. Shaded nodes represent observed variables, empty nodes represent latent variables, and striped nodes represent latent variables inferred prior to running the model. The *K* plate contains loadings for topics shared across modalities, while the *K^s^* plate surrounds genotype-specific topics.

In this multi-modal version of LDA, the same latent factors are shared across data modalities, allowing features of each modality to be directly linked together. We include a modality-specific (private) subspace for genotype, *βτ*, to control for ancestral structure in mixed-population samples. The weight of the private versus shared genotype subspaces is determined by 0 < *α* < 1. The model does not include a gene expression-specific latent space to avoid losing any broad regulatory signal that is genotype-dependent (Rakitsch and Stegle 2016), as is often the case with trans-eQTLs (GTEx Consortium 2017). Critically, because our framework directly models count data, we avoid spurious or distorted signals through data normalization (Hicks et al. 2018; Love et al. 2014; Robinson et al. 2010), and due to the non-negative factors, the components capture parts-based patterns instead of global patterns (Lee and Seung 1999; Townes and Engelhardt 2021). The multinomial distribution allows us to separate out variation due to library-size effects from the underlying compositional variation, which is more biologically-relevant.

Features that have higher weights within a topic 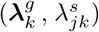 have a larger relative contribution. However, the proportion of total counts for each gene varies widely. Genes with higher counts may dominate certain topics merely due to their high expression levels, overshadowing lower-expressed genes that are actually more informative for that topic compared to others. Consequently, instead of using the raw expected loadings, we determine the importance of each feature across topics by ranking the average 2-Wasserstein distance between the posterior variational distributions. This allows us to control for both average feature counts and varying uncertainty in model estimation by using the full information provided by the posterior estimates. SNP minor allele frequency (MAF) is much less variable than gene total counts. To control for allele counts, we rank SNPs after regressing out the coded MAF in each loading (see Methods for details).

## Results

We applied our telescoping bimodal LDA (TBLDA) model to gene expression data for the ten tissues with the highest number of samples from the v8 GTEx data release, and to the genotypes from all individuals who contributed to at least one of the samples (Table 1). We took advantage of the known GTEx covariates to interpret biological variation captured in the model factors and ensure relevant signal was found.

**Table 1.** Overall summary statistics and available data for each of the ten tissues included in the analysis.

| Tissue | Sample Size | Num. Tissue-Associated Factors | Cell Type Enrichment Scores |
| --- | --- | --- | --- |
| Subcutaneous Adipose | 581 | 1,536 | Adipocytes |
| Tibial Artery | 584 | 1,504 |  |
| Esophagus Mucosa | 497 | 1,254 | Keratinocytes, Epithelial Cells |
| Lung | 515 | 1,095 | Epithelial Cells |
| Skeletal Muscle | 706 | 2,649 | Myocytes |
| Tibial Nerve | 532 | 905 |  |
| Skin (Not Sun-exposed) | 517 | 1,225 | Keratinocytes, Epithelial Cells |
| Skin (Sun-exposed) | 605 | 1,390 | Keratinocytes, Epithelial Cells |
| Thyroid | 574 | 1,282 | Epithelial Cells |
| Whole Blood | 670 | 2,599 | Neutrophils |

First, we checked that ancestral structure, using reported ancestry as a proxy, was not associated with the estimated shared factors. As expected, ancestral structure was largely controlled for since it is captured in the genotype-specific portion of the model (median absolute value factor-ancestry point biserial correlation coefficient < 0.01).

Next, we looked for signal from one of the top sources of known variation in the dataset, tissue of origin, by identifying factors active in specific tissues. We found 15,439 tissue-factor associations via the inner product of each factor and a tissue indicator vector, considering inner products greater than 40 to be tissue-associated (Fig. S2, Table 1). Tissue sample size was strongly correlated with the number of tissue-associated factors (Kendall’s rank correlation *τ* = 0.64, *p* < 0.01), which suggests that certain tissues may have underpowered downstream analyses. Over all runs, whole blood and skeletal muscle had the most associated factors (2, 599 and 2, 649 respectively) while tibial nerve had the fewest (905). Skin (sun-exposed), skin (not sun-exposed), lung, subcutaneous adipose, and thyroid had the weakest associations (median inner products between 60.7 and 69.7); whole blood, tibial nerve, and esophagus mucosa have the strongest (median inner products between 88.6 and 109.6). Accordingly, whole blood and skeletal muscle samples allocate a majority of their topic membership into tissue-specific factors (Fig. 2).

**Figure 2.**
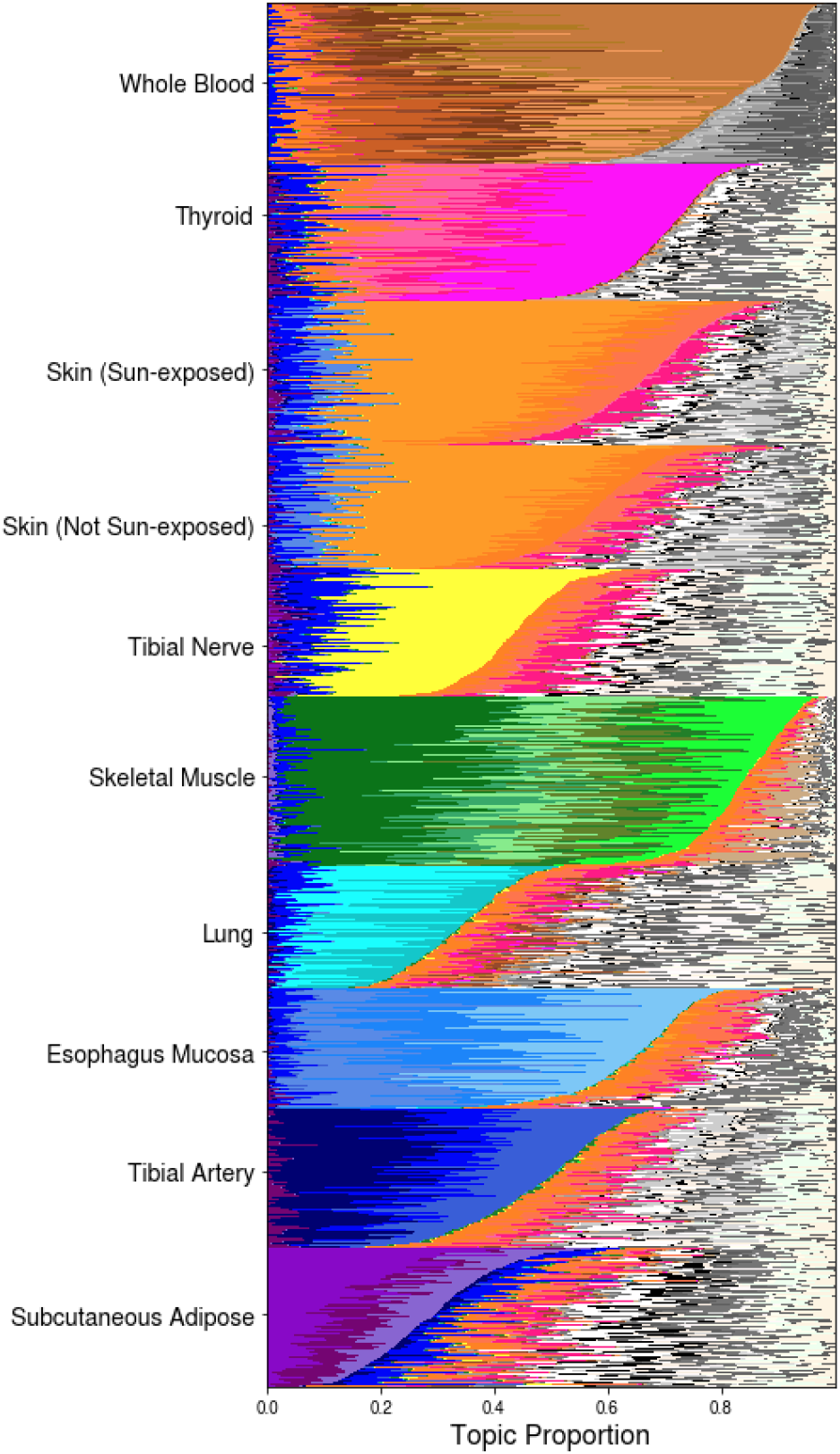
The estimated TBLDA topics capture tissue-specific signal. Each row depicts the expected sample-topic proportion (*ϕ_l_*, eqn. 2) for one sample for the model fit using genes on chromosome 19 and SNPs on chromosome 22; samples are sorted by tissue. Tissue-associated topics are colored in a tissue-wise family color scheme. The remaining topics are drawn using a random gray scale.

In order to explore the robustness of these tissue-associated factors, we compiled sets of the top-ranked features that frequently appeared across factors associated with a common tissue (see Methods). Taken together, the two skin tissues had the largest group with 6,515 genes, while whole blood had the largest number of unique genes considering the other sets (1,983). All tissue-associated robust gene sets were enriched for functionally-relevant biological process Gene Ontology (GO) sets (Benjamini Hochberg (BH) FDR < 0.1, Table S1). Further, all relevant robust tissue gene sets (except for tibial artery) contained a majority of the tissue-specific transcription factors (TFs) present in the overall analysis (whole blood 10/12, thyroid 6/8, esophagus 7/8, skins 12/14, lung 3/4, nerve 3/3, skeletal muscle 12/12, tibial artery 0/2, subcutaneous adipose 2/2; see Methods). This demonstrates that our model consistently found topics that captured important tissue-specific biological variation including functional pathways and tissue-specific regulatory activity.

Next, we used a compilation of 63 SNP classes from the LDSC (Bulik-Sullivan et al. 2015) data repository to explore functional regulatory enrichments among tissue-associated SNPs. The union of all robust tissue-associated SNPs was enriched for 20 SNP classes (Fisher’s exact test, BH FDR < 0.1) including TFBS_ENCODE (BH FDR < 0.016), SuperEnhancer_Hnisz (BH FDR ≤ 3.4 × 10^−3^) and active enhancer-associated H3K27ac_Hnisz (BH FDR ≤ 9.8 × 10^−5^) and H3K4me1_Trynka (BH FDR ≤ 2.6 × 10^−5^). In particular, several single tissue-associated SNP sets are associated with DGF_ENCODE, DHS_Trynka.extend.500, H3K4me1_Trynka, H3K27ac_Hnisz, and Enhancer_Hoffman (eight, six, five, five, and three tissues respectively out of the ten total; BH FDR < 0.1; Table S2). These SNP set enrichments from our model show that TBLDA identifies functional connections between genotype and gene transcription; these enrichments are intriguing because trans-eQTLs are known to be associated with enhancer activity (GTEx Consortium 2017).

We then investigated whether the SNP sets are clustered together in particular genomic regions. The union of all tissue-associated SNPs was not enriched in any chromosomal regions using a bin size of 250,000 bp and a sliding window of 100,000 bp, but there were 143 tissue-specific genomic bin enrichments (Fisher’s exact test, FDR < 0.1; Fig. 3, Table S3). Notably, 48 regions on chromosome four were enriched for the robust SNP set associated with subcutaneous adipose (BH FDR < 0.05). This highlights the ability of TBLDA to identify jointly functional genomic regions even when the SNP data have been LD-pruned.

**Figure 3.**
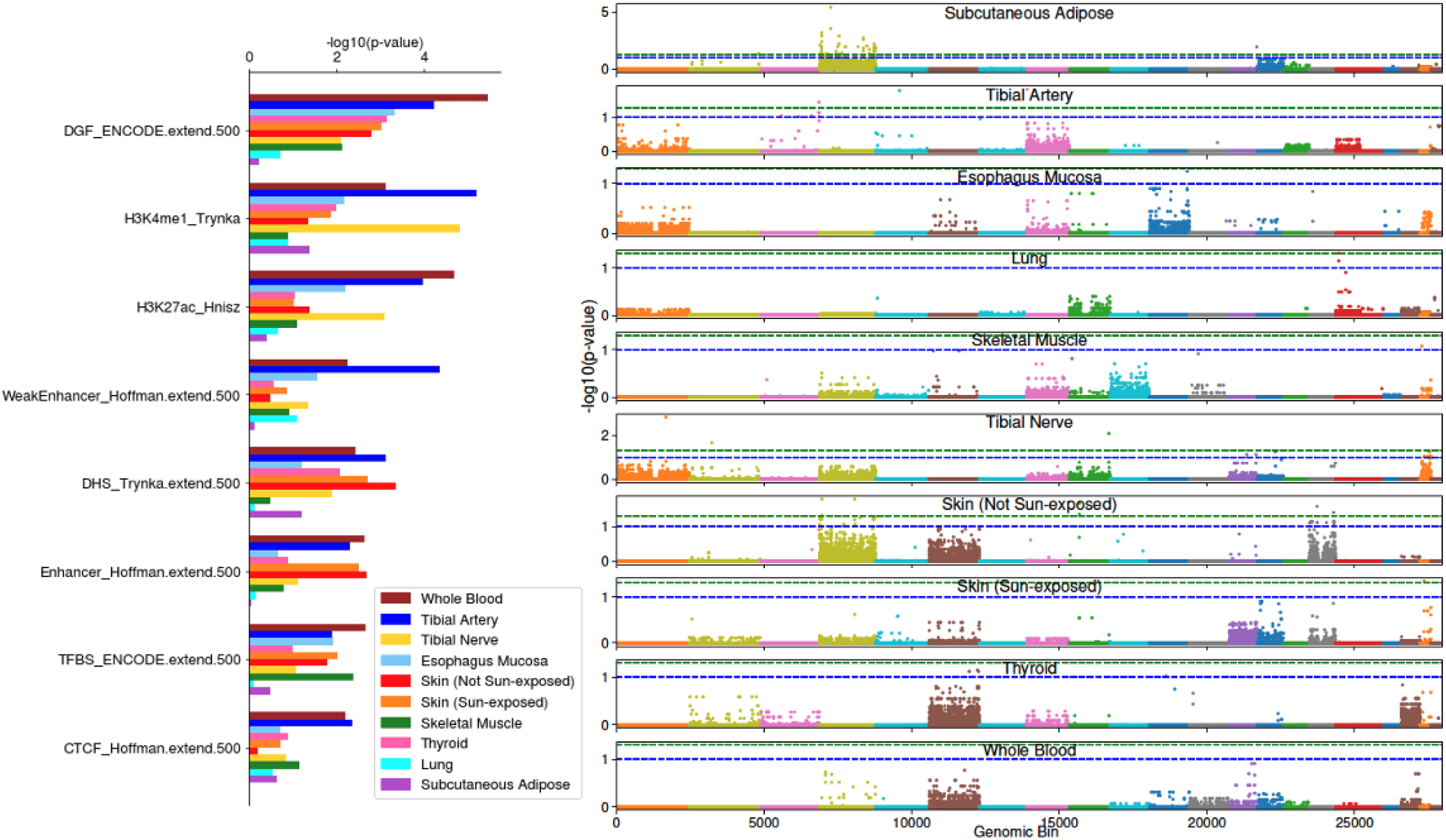
Robust tissue-associated SNP sets are enriched for DNA markers and localized throughout the genome. Left: Enrichment via Fisher’s exact test of eight of the 63 LDSC SNP classes across all robust tissue-associated SNP sets. Right: Each tissue’s associated SNP set was tested for genomic localization via Fisher’s exact test. The blue and green dotted lines are drawn at p-value thresholds of 0.1 and 0.05 respectively. The colors mark the division between ordered chromosomes, with chromosome one on the far left.

Although the GTEx data provide the ground truth of each sample’s origin tissue, this is not the case across all data sets. Thus, we next evaluated whether our model could recover robust components across relevant runs in an unsupervised manner. To do this, we ran our model 484 times, once for each pair of chromosomes in the GTEx v8 data, and identified shared components across these runs (see Methods). Across all runs, we recovered 197 clusters of robust genotype factors and 1,799 groups of robust gene expression factors. Loadings that were well-correlated with each other across runs tended to cluster by tissue; 81 of the robust genotype clusters and 468 of the robust gene expression clusters included factors that were associated with the same tissue (Fig. 4). Only 14 and 75 of the robust genotype and expression clusters, respectively, did not include tissue-associated factors. The presence of these tissue-associated robust genotype components demonstrates that TBLDA identifies interactions between the data modalities versus separate structure within each modality.

**Figure 4.**
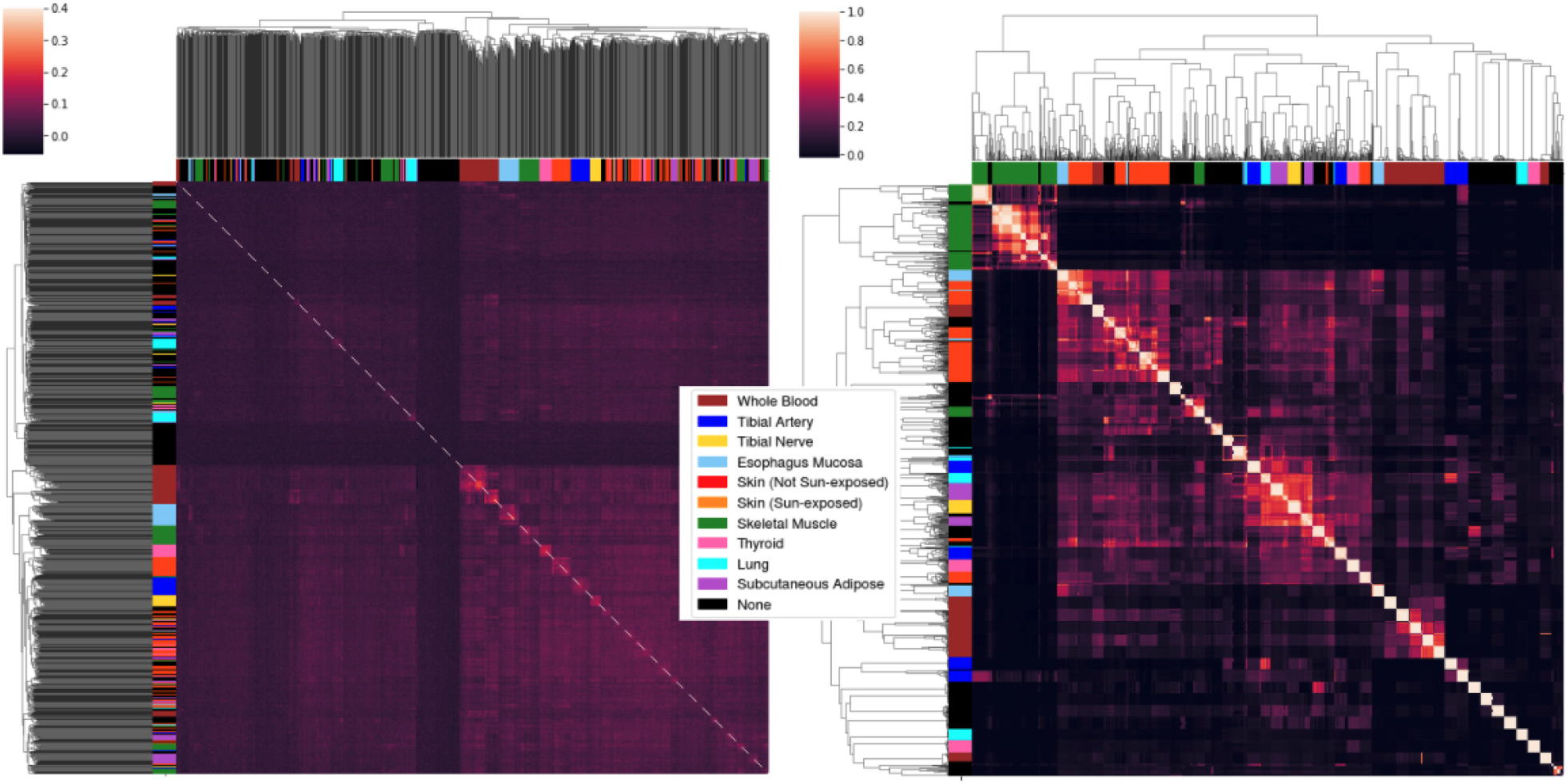
The TBLDA model estimates robust factors across independent runs. Cluster maps of the pairwise Pearson correlations between loadings from all runs that used features from chromosome two. The color bars associated with the axes label the topic’s strongest tissue association, if any. Left: Correlations calculated using the residuals after regressing coded MAF out from the expectation of the genotype loadings 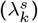. Right: Correlations between the expected value of gene expression loadings 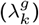.

Due to the nature of bulk RNA-seq expression data, the GTEx samples average expression over heterogeneous tissue samples containing various cell types. We computed the Kendall correlation between cell-type enrichment scores and factor values to determine whether factors capture sample cell-type composition (Fig. S3). We use estimated enrichment scores for bulk cell deconvolution across five cell types (adipocytes, keratinocytes, epithelial cells, myocytes, and neutrophils) in eight tissues for a total of 11 tissue- and cell-type pairs (Table 1) (GTEx Consortium et al. 2020). Enrichment scores for 8/11 pairs of tissue and cell-types were well captured by at least one factor (maximum abs(Kendall *τ*) > 0.5). This suggests that the TBLDA components often represent cell-type specific processes within tissue samples.

To test whether traditional eQTLs ascertained using univariate tests are captured by TBLDA, we ran a linear model for association between the top 10% most informative SNPs and genes on common factors for each tissue separately using MatrixEQTL (Shabalin 2012) (further referred to as the multivariate testing approach; Fig. 5). Out of 10,855,277 total tests, we found 53,358 cis-eQTLs at BH FDR < 0.1 across all ten tissues including 14,971 unique eVariants and 8,913 unique eGenes (Table S4). The majority of these cis-eQTLs (46,351) affect protein-coding genes, with a minority (7,007) acting on linc-RNA genes (Fig. 5). Thyroid had the most cis-eQTLs, 6,590, followed by tibial nerve with 6,182.

**Figure 5.**
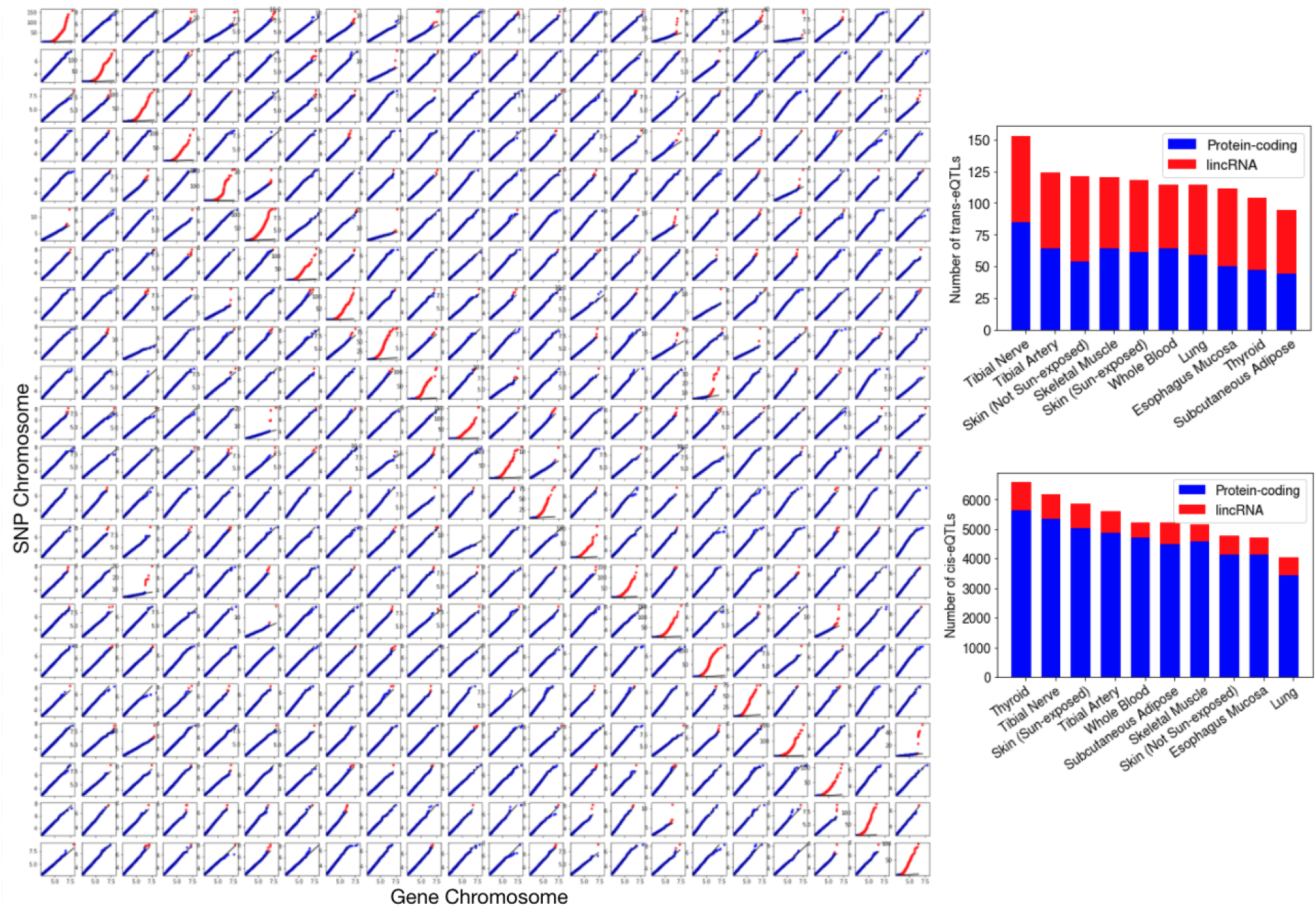
Characterization of cis- and trans-eQTLs between top-ranked features in each factor. Left: For each of the 484 model runs, the ordered true MatrixEQTL association −log10(p-values) (y-axis) are plotted against ordered −log10(p-values) from tests using the same features but permuted expression and covariate data (x-axis). Clear cis-eQTL enrichment is present across all intrachromosomal runs. Points at which the ordered true −log10(p-value) is greater than the maximum permuted −log10(p-value) are colored in red to highlight deviation. Right: Histograms depicting the numbers of trans- (top) and cis- (bottom) eQTLs mapped per-tissue, split by gene type.

We also discovered 1,173 trans-eQTLs at BH FDR < 0.1, which include 961 unique trans-eGenes and 1,074 unique trans-eVariants (Table S4). In contrast to both the data, which consist of 86% protein-coding genes, and the cis-eQTLs, trans-eQTLs have an almost equal number of linc-RNA and protein-coding eGenes (581 and 592, respectively; Fig. 5). Tibial nerve had the highest number of trans-eQTLs (153), followed by tibial artery with 124. Not surprisingly (GTEx Consortium 2017), the cis-eQTL enrichment is much stronger and more consistent than the trans-eQTL enrichment (Fig. 5). These eQTL mapping results highlight the associations between SNPs and genes loaded onto a common factor, and suggest that eQTL candidates may be identified using the TBLDA factors.

Next, we restricted our analysis within each tissue to the respective tissue’s associated factors. We found 7,968 cis-eQTLs (4,282 unique eGenes, 5,922 unique eVariants) and 1,081 trans-eQTLs (983 unique trans-eGenes, 1,073 unique trans-eVariants) at BH FDR < 0.1 (Table S5). Whole blood had the most cis-eQTLs while skin (not sun-exposed) had the most trans-eQTLs. Similarly to the full analysis above, the trans-eQTLs have an approximately equal split of linc-RNA and protein-coding eGenes whereas the cis-eGenes are mostly protein-coding (Fig. S4). Of the discoveries, 962 (89.0%) of the trans-eQTLs and 2,025 (25.4%) of the cis-eQTLs were novel, meaning not below significance threshold in the unrestricted multivariate test. The fact that 1,054 (89.9)% of the trans-eQTLs found by the full multivariate test were not found in the tissue-associated factors indicates that most of the trans associations found by the model are not in specific tissue-factor pairs.

Thus, to increase power to find trans-eQTLs shared across tissues, we next limited association tests to features in general factors that were not linked to any tissue. This approach yielded 1,065 trans-eQTLs (895 unique trans-eGenes, 1,007 unique trans-eVariants) and 23,965 cis-eQTLs (5,111 unique eGenes, 7,229 unique eVariants; Fig. S4, Table S6). Here, skin (sun-exposed) had the most trans-eQTLs (124) while thyroid produced the most cis-eQTLs (2,883). The reduction in test numbers allowed 578 new trans-eQTLs and 2,393 cis-eQTLs to move below the significance threshold relative to the unrestricted multivariate test.

Inferred covariates such as PEER factors are known to capture and thus inadvertently control for broad regulatory effects that potentially have a true genetic basis, potentially removing broad trans-eQTL signals (GTEx Consortium 2017; Rakitsch and Stegle 2016). To test whether factors in our model find these kinds of regulatory hotspots, we ran the same eQTL mapping as before except excluding all PEER factors from the covariate matrix. This resulted in fewer total cis- and trans-eQTLs (16,731 and 1,004, respectively at BH FDR < 0.1; Table S7). However, the proportion of unique eVariants to trans-eQTLs versus including PEER factors was lower (0.80 versus 0.92), suggesting that, to some extent, PEER factors do remove trans-acting pleiotropic signals that are captured by our model. In line with their supposed mechanisms of action, 95.7% (16,019) of these cis-eQTLs overlapped with our prior analysis controlling for PEER factors, while only 11.0% (110) of these trans-eQTLs were also found when controlling for PEER factors in the association analysis.

Next, we explored the overlap of our eQTLs and the GTEx consortium cis- and trans-eQTL list, produced by the consortium through an exhaustive tissue-specific testing approach (GTEx Consortium et al. 2020). All except for three of the multivariate TBLDA cis-eQTLs were in the GTEx cis-eQTL list. Of the multivariate trans-eQTLs, 15 overlapped with the 2,629 genome-wide GTEx trans-eQTLs in the top ten tissues. However, 712 (27.1%) of the GTEx trans-eQTLs included one of those 15 eGenes. That number is largely driven by four eGenes in three tissues that are associated with hundreds of SNPs in an LD block with the relevant eVariant in our analysis. Notably, 284/428 (66.4%) of lung trans-eQTLs had *LAMA2* as a trans-eGene, 176/795 (22.1%) of thyroid trans-eQTLs act on the protein-coding gene TMEM253, 59/439 (13.4%) of skeletal muscle trans-eQTLs involve the protein-coding gene *PARP10*, and 97/795 (12.2%) of thyroid trans-eQTLs have *MAPRE3* as a trans-eGene. Although we fail to capture this extended signal because we use an LD-pruned SNP set, our model still groups these genes and their genomic hotspots together. To more directly evaluate the impact of LD-pruning, we looked for GTEx trans-eQTLs that matched ours in tissue and eGene but included an eVariant within 100 Kb upstream or downstream of ours. We found that 30.8% (810) of the GTEx trans-eQTLs overlap ours in terms of this LD-proxy test. Taken together, these results suggest that our approach finds overlapping cis-eQTL signals but expands our ability to identify broad-acting trans-eQTLs in these bulk data.

One interesting example is the trans eVariant rs4297160, which is associated with both *MAPRE3* (p-value p ≤ 6.6 × 10^−11^) and *ARFGEF3* (p-value p ≤ 2.2 × 10^−16^) in thyroid and sits in the 9q22 locus (Fig. 6). Specifically, rs4297160 is located within the lincRNA gene *PTCSC2*, which has been linked to a predisposition for papillary thyroid cancer (He et al. 2015). The 9q22 locus, home of thyroid-specific TF *FOXE1*, was previously found associated in trans with *ARFGEF3* in thyroid (GTEx Consortium 2017). Notably, PEER factors were shown to capture and therefore control for broad regulatory signals from that locus (GTEx Consortium 2017); in line with this, in our association tests run without PEER factors as covariates, rs4297160 was a trans-eVariant for 34 different genes in thyroid, including *HECW1* (regulates the degradation of thyroid transcription factor 1 (Liu et al. 2019)), *COL-GALT2* (downregulated in patients with thyroid orbitopathy (Khong et al. 2015)), and *FMO5* (expressed, in endocrine cells that produce hormones that regulate metabolism (Xu 2017)); Fig. 6).

**Figure 6.**
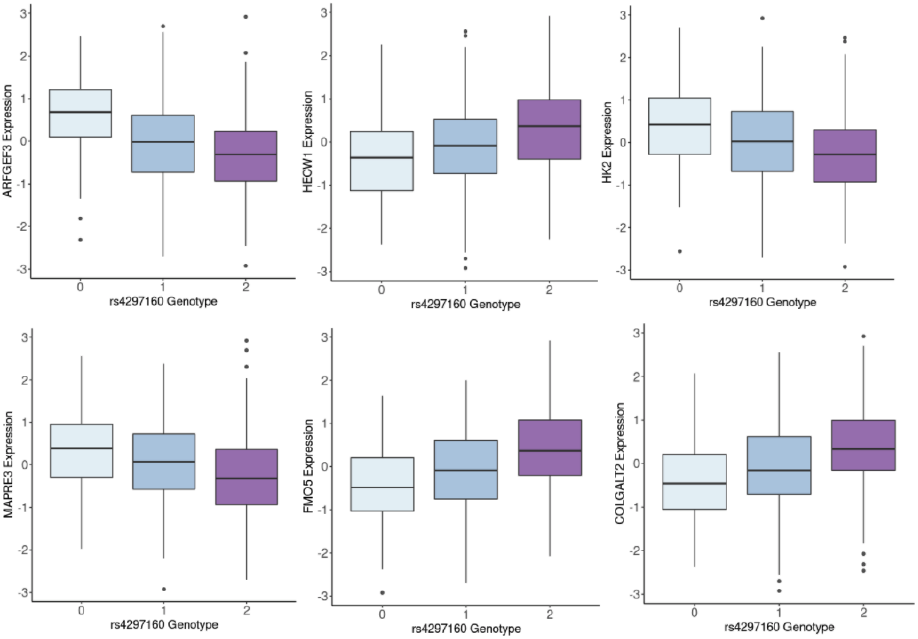
Example trans-eQTLs stemming from a single locus in thyroid. Gene expression counts from thyroid samples have been quantile normalized.

We evaluated the increase in statistical power compared to the univariate approach due to our reduced multiple testing burden. The cis-eQTL p-values with BH FDR < 0.1 from our method have a different distribution from the GTEx cis-eQTLs found via exhaustive search (Kolmogorov-Smirnov test statistic 0.37, p-value *p ≤* 2.8 × 10^−14^). Although our cis-eQTLs found via TBLDA are a subset of all true associations, they tend to have stronger associations than the set of GTEx cis-eQTLs (Fig. 7).

**Figure 7.**
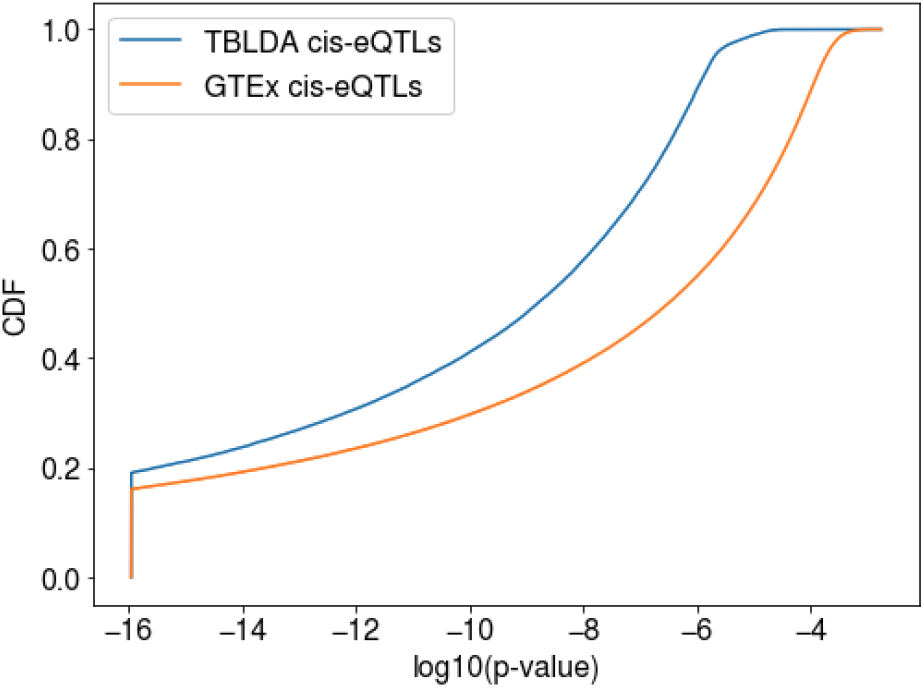
Empirical CDF comparison between our cis-eQTL p-values and those found via the exhaustive GTEx search. Cis-eQTL p-values are trimmed to double machine precision, and then used for the Kolmogorov-Smirnov test.

## Discussion

In this paper, we present a probabilistic telescoping bimodal latent Dirichlet allocation (TBLDA) model that uncovers shared latent factors between bulk RNA-seq expression and genotype data when there is not a one-to-one mapping among the samples for each data modality. The model takes raw counts as input, which avoids any potential data skewing due to normalization. We fit the model using gene expression data from ten tissues in the GTEx v8 release and matched donor genotypes. Using known GTEx covariates, we established that the recovered topics reflected meaningful biology such as sample cell-type proportion (Fig. S3). Robust gene sets in tissue-associated factors were enriched for functionally-relevant pathways (Table S1). Causal eVariants identified by our method are known to be enriched in a variety of genomic regulatory regions (Albert and Kruglyak 2015); top-ranked robust tissue-associated SNP sets in our model were likewise enriched, demonstrating motifs of known eVariants (Table S2).

Running linear association tests on top-ranked features from each factor using MatrixEQTL (Shabalin 2012), we found 53,358 cis-eQTLs and 1,173 trans-eQTLs at BH FDR 0.1, including four trans-eGenes that accounted for 66%, 22%, 13%, and 12% of the trans-eQTLs in their respective tissues in the GTEx v8 trans-eQTL analysis. By restricting association tests to the top features per factor in our model, we decrease the multiple testing burden and increase power for mapping trans-eQTLs. This is demonstrated by the fact that 1,158 of our trans-eQTLs were not identified in the exhaustive genome-wide GTEx analysis. A critical caveat of our approach is that, with a finite number of topics, we do not expect the model to capture all true eQTLs; however, we show that it does reproducibly identify novel and functionally-relevant eQTLs. Taken together, though, these results demonstrate that our method successfully learns biologically meaningful shared topics across gene expression and genotype data.

There are several potential points of contention in our model. First, although the model’s probabilistic nature provides important measures of uncertainty for noisy genomic data, due to our inference procedure the posterior should be interpreted with caution since variational inference is known to underestimate the posterior variance (Giordano and Broderick 2015). Second, because we do not include a private subspace for gene expression, true latent components that reflect expression-specific variation such as batch effects will be forced to contribute to the modality-shared factors. We believe this is important in order to retain signal for broad regulatory effects that especially affect trans-eQTL discovery. Nevertheless, if the model is used in a context such as single-cell RNA-sequencing, where there are known and strong expression-specific covariates such as batch effects, this design choice should be reconsidered. Furthermore, a natural question that arises for all parametric latent factor models is how to determine the number of topics. We stress that there is no ‘correct’ topic number, and the user will want to make a reasonable trade-off between computational speed for inference and the granularity of signal captured. In practice, we recommend anywhere from 20 to 150 factors depending on the size of the data set. Given these qualities, natural extensions to the model include adding latent or semi-supervised expression-specific topics and extending it to a nonparametric framework.

## Extended Methods

### Feature Selection

Following (Jo et al. 2016), we used plink 1.9 (Purcell et al. 2007) to trim the GTEx v8 Whole Genome Sequencing SNP sets such that no two SNPs within a 200 Kb window have a Pearson correlation ≥ 0.2. SNPs with imputed genotypes were removed, yielding 202,111 remaining SNPs across 831 individuals. All gencode v26 autosomal lincRNA and protein-coding genes from the 5,781 samples with genotypes were considered. We retained the 19,534 autosomal genes with a median RNASeQC v1.1.9 (Graubert et al. 2021) read count of at least five in at least one tissue. SNPs and genes were split into 22 groups by chromosome.

### Ancestry Structure

We ran terastructure (Gopalan et al. 2016) on the 202,111 LD-trimmed SNPs with the following options: `-rfreq=40222 and −K=5`. The resulting beta and theta output matrices were assigned to *β* and *τ* (eqn. 8) to produce the genotype-specific portion of the model.

### Model Runs

We used Pyro v1.4.0 (Bingham et al. 2019)’s stochastic variational inference framework to fit the model, using `pyro.poutine.scale`(scale=1.0 × 10^−6^) for numerical stability, an Adam optimizer, and a learning rate of 0.05. ξ and *σ_i_* were set to one, *δ* to 0.05, and *μ* to 0.85. The model was fit separately for feature sets from each chromosome combination, for a total of 22 × 22 = 484 runs. Let *x* be the average ELBO over the latest 1000 epochs and *y* be the average ELBO over the 1000 epochs prior to those. Runs were terminated when 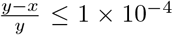. Code to run TBLDA is available at https://github.com/gewirtz/TBLDA.

### Feature Ranking

After regressing out allele frequency, we take the top 10% of features from each loading with the highest absolute value residuals. The 10% of genes from each loading with the highest 2-Wassterstein distances are considered the top gene features. Because the feature numbers vary by chromosome, runs have differing numbers of top features associated with their factors.

### Functional Enrichment Data

The tissue-specific TF list originated from Table S3 in (Sonawane et al. 2017). To conduct GSEA, we used all biological process terms from GO v6.2 that had at least three genes in common with our analysis feature set. We used LDSC’s baselineLD v2.1 (Bulik-Sullivan et al. 2015; 2021) genome annotations to compute SNP set enrichments. We did not consider MAF_bin classes.

### Tissue-Associated SNPs

We computed the inner product between tissue indicator vectors and the expectation of the variational posterior for phi. The top-ranked SNPs in topics with an inner product > 40 were considered associated with a specific tissue within each run. For each tissue, we calculated the 75th percentile of the distribution of total tissue-associated factors across all runs that each top-ranked SNP is associated with. SNPs that are top-ranked in at least the 75th percentile of each tissue’s associated factors across all runs comprise the set of robust tissue-associated SNPs.

### eQTL Pipeline

We used MatrixEQTL v2.3 (Shabalin 2012) with modelLINEAR to run the eQTL testing. Expression for all genes that passed a 0.8 mappability filter was quantile-normalized as input. Sex, PCR, platform, the top five genotype principal components, and the top 60 PEER factors per tissue were included as covariates. FDR was computed using the Benjamini-Hochberg procedure over each run for protein-coding and lincRNA genes separately.

### Robust Components

We computed the correlation of each factor loading with all other loadings from runs on the same chromosome. Any factor with more than two loading Kendall *τ* > 0.15 for SNPs and three Pearson *r*^2^ > 0.95 for genes was flagged—along with the highly-correlated factors—as a robust component. For each robust component, we averaged the constituent loadings to produce a representative factor loading. All components whose representative loadings exceeded *r*^2^ > 0.95 were further collapsed into a single robust component.

### Data Access

All raw sequencing and genotype data from GTEx v8 used in this study can be found in dbGaP under accession number phs000424.v8.p2.

## Author’s contributions

AG, FWT, and BEE designed the method. AG implemented the method and conducted data analysis. AG and BEE wrote the manuscript. AG, FWT, and BEE edited the manuscript.

## Acknowledgements

AG, FWT, and BEE were funded by Helmsley Trust grant AWD1006624, NIH NCI 5U2CCA233195, NIH NHLBI R01 HL133218, and NSF CAREER AWD1005627.

## Additional Files

Additional file 5 — Supplemental Table S1. Gene set enrichment analysis of tissue-associated robust gene sets.

All KEGG, Biocarta, GO biological process, and Reactome gene sets with BH FDR < 0.1 in at least one tissue-associated robust gene set.

**Figure 8 Supplemental Figure S1.**
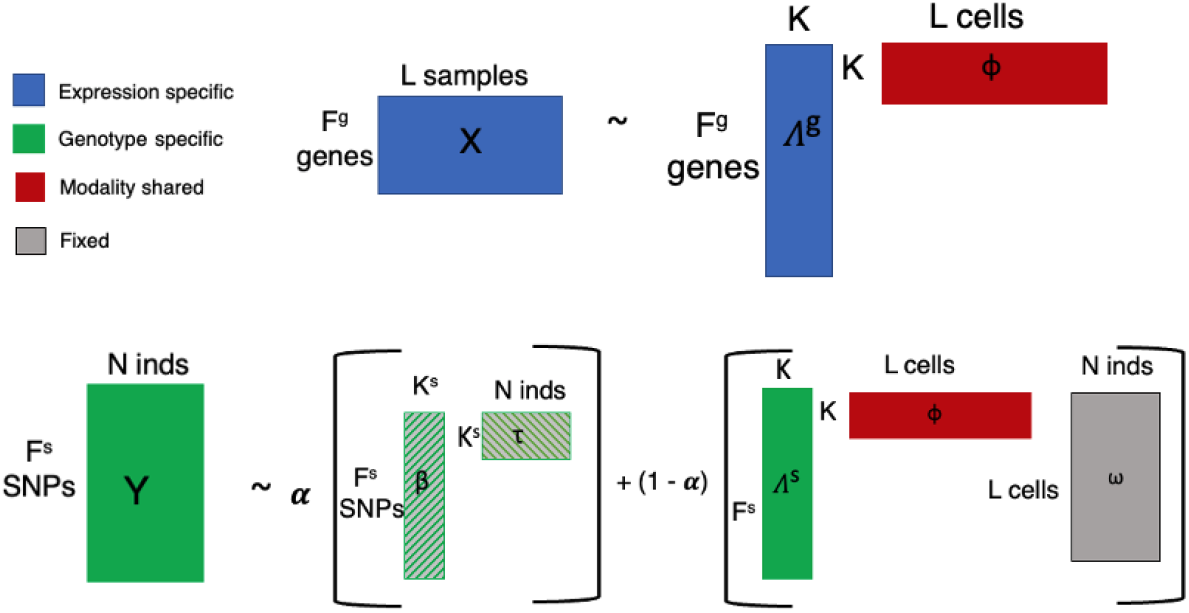
Model visualization. Explicit representation of the model with dimensions drawn out. Each portion of the model is color-coded according to modality. Gray represents the known mapping between samples and individuals. The ancestry portion is striped because while it is learned, that occurs prior to fitting the shared model portion.

Additional file 6 — Supplemental Table S2. LDSC SNP class enrichment among tissue-associated robust SNP sets.

Additional file 7 — Supplemental Table S3. Genomic regions enriched for tissue-associated robust SNP sets (BH FDR < 0.1).

Additional file 8 — Supplemental Table S4. Cis- and trans-eQTLs mapped using the multivariate testing approach (BH FDR < 0.1).

Additional file 9 — Supplemental Table S5. Cis- and trans-eQTLs mapped using only tissue-associated topics (BH FDR < 0.1).

Additional file 10 — Supplemental Table S6. Cis- and trans-eQTLs mapped using general topics (not associated to any tissue; BH FDR < 0.1).

Additional file 11 — Supplemental Table S7. Cis- and trans-eQTLs mapped using the multivariate testing approach, without including PEER factors as covariates (BH FDR < 0.1).

**Figure 9 Supplemental Figure S2.**
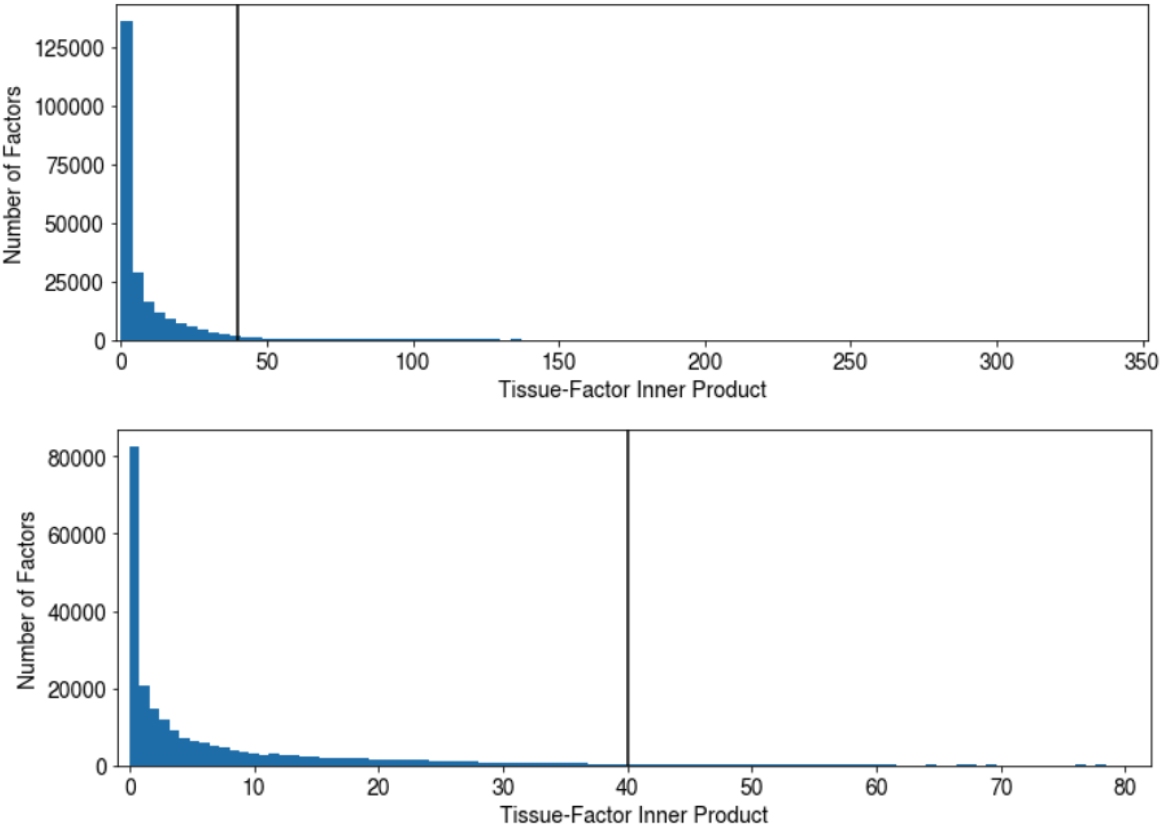
Histogram of tissue-factor inner products. The vertical line depicts the inner product threshold for determining tissue association. The bottom figure presents the same data as the top figure, but zoomed in to lower counts to illustrate the long tail. Both histograms are drawn using 100 bins.

**Figure 10 Supplemental Figure S3.**
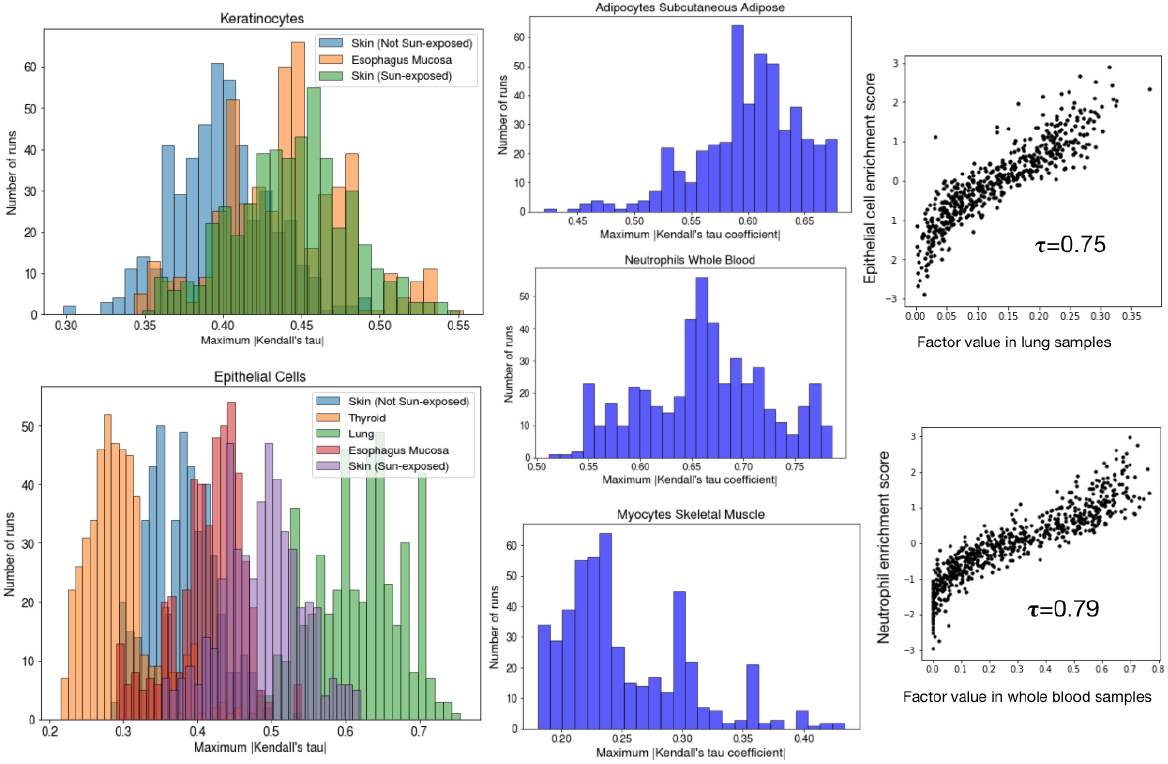
Relationships between factors and cell type enrichment scores. Left and center: Histograms depicting the maximum absolute value Kendall’s tau coefficient in each of the 11 available tissue-cell type pairs (Table 1) Right: Scatter plots of the expected factor values (*ϕ_l_k*) and cell type enrichment scores for two strongly correlated examples.

**Figure 11 Supplemental Figure S4.**
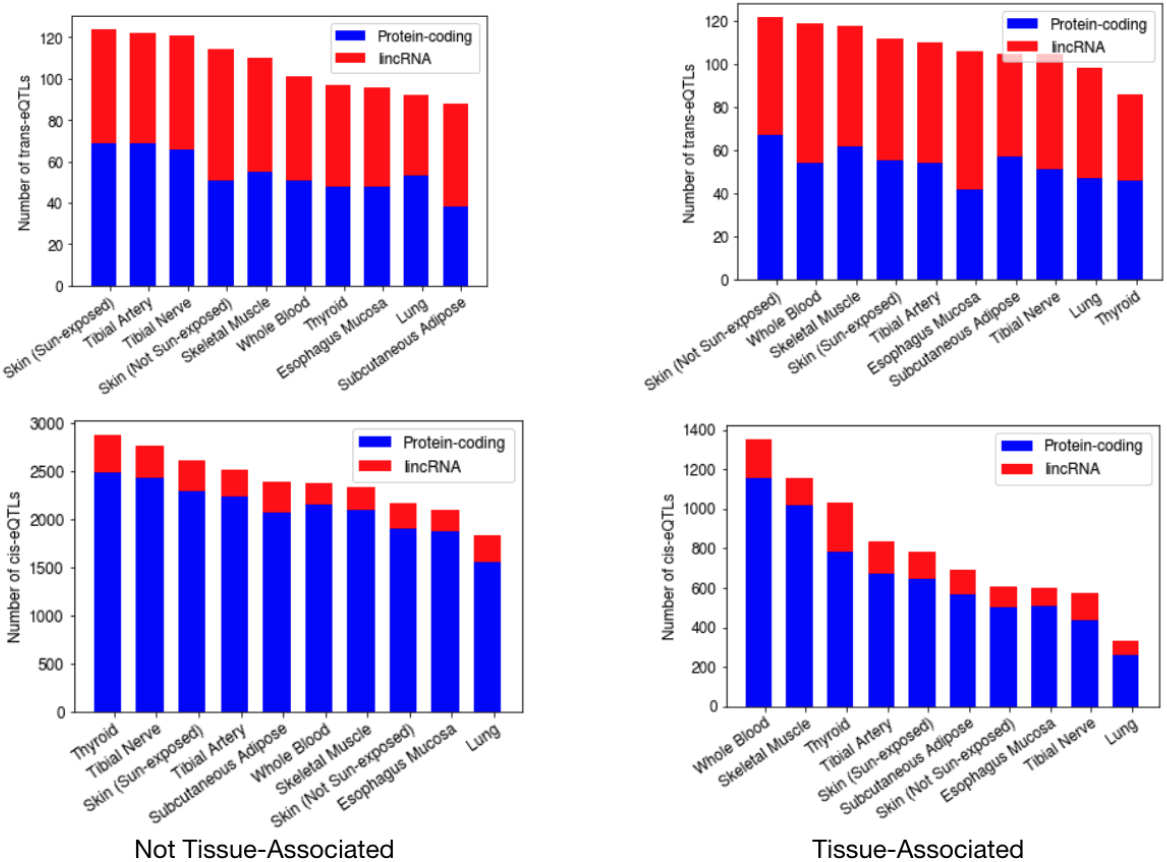
Significant eQTL counts from restricted tests. Per-tissue, the number of eQTLs found when limiting association tests by using only non-tissue-associated factors (left) and restricting by tissue with only tissue-associated factors (right).

